# Optimizing Electrotactile Stimulation Parameters for Responses under Cognitive Load

**DOI:** 10.1101/2025.09.08.674827

**Authors:** Felix Jarto, Elaine Corbett, Sigrid Dupan

## Abstract

**Background:** Prosthetic users mainly rely on their vision for feedback during control of the device in absence of the tactile and proprioceptive modalities. Overreliance on vision increases cognitive load during prosthesis use, resulting in reduced control performance. Provision of additional feedback through other modalities could reduce this overreliance on vision and improve closed-loop prosthetic control. Fast and accurate recognition of feedback is particularly important when we consider that both delays and inaccuracies can destabilize closed-loop control environments. Transcutaneous electrotactile stimulation is a promising technology with advantages including non-invasiveness and simplicity. It is not yet clear, however, which parameters influence the speed and accuracy of response to electrotactile stimuli. In this study, we set out to investigate how we can influence response characteristics to an instantaneous change in stimulation intensity by manipulating both stimulus-related and environmental variables.

**Methods:** 20 participants completed a randomized reaction time test to both visual and electrotactile stimulation. Participants were asked to respond to incoming stimuli with a button press on a controller as fast as possible, without discriminating between modality of stimulation. In a second experiment, 20 participants completed an intensity discrimination task for electrotactile stimulation. Participants had to prioritize either speed or accuracy during specific blocks, while cognitive load, magnitude of intensity shift and direction of shift were manipulated.

**Results:** The results of the reaction time experiment confirmed lower average response times for electrotactile stimuli compared to visual stimuli by a median of ~50ms (p=0.008). In the intensity discrimination experiment, increased shift intensities led to increased response accuracies and decreased response times (p<0.001 in both cases). The presence of cognitive load increased average response times (p=0.005), but did not affect response accuracies (p=0.420).

**Conclusions:** The results of the intensity discrimination experiment imply that the magnitude of intensity shift needs to be provided via steps multiple times larger than the just noticeable difference to ensure fast and accurate responses to electrotactile stimulation. Moreover, results from the reaction time experiment confirmed faster average response times for electrotactile stimuli over visual stimuli. We argue that providing electrotactile stimulation using bigger steps in perceived intensity could result in the supplementary feedback making closed-loop control of prosthetic devices more reliable.

## Introduction

In the last two decades upper limb prosthetics have seen many technological improvements, yet rejection rates continue to remain at levels as high as 44%^[1]^. Previous studies have shown that major reasons behind device abandonment include, among others, unreliable control and lack of sensory feedback^[2,3]^. The topic of sensory feedback - both invasive and non-invasive - has been heavily investigated in recent years: many studies have found promising results regarding enhanced functionality^[4]^, promotion of embodiment^[5]^, decrease in cognitive load^[6]^ and alleviation of phantom limb pain^[7]^. However, it is difficult to draw conclusions, as currently there is no gold standard for sensory feedback and many different feedback strategies are employed in the aforementioned studies. Moreover, quality-of-life improvements outside of the lab have not been established yet. There is a lack of data collected in home environments^[8–10]^, and those that collected data at home reported significant variabilities in the performances of their subjects^[11]^. Sensinger and Dosen hypothesize that mixed outcomes and, by extension, the lack of sensory feedback in commercially available prostheses are a result of insufficient understanding about the role and mechanisms of feedback in prosthetic control^[12]^. Thus, to improve sensory feedback paradigms for prosthetics, it is crucial to investigate sensory feedback through the lens of sensorimotor control.

The overall accuracy of our movements is amazing, especially if we take into account that both our body and the environment change over time. However, when studying movement in more detail, it is clear that their outcomes do not always align with our intentions. One contributing factor to this misalignment is inherent noise in the sensory and motor systems, mainly related to synaptic activity^[13]^. Another important factor is that movement is based on internal models representing our body and the environment^[14]^. When changes happen, e.g. when our muscles are fatigued, misalignment will occur. When the executed movement doesn’t align with our intentions, sensory feedback allows us to perceive the difference, make small corrections on the fly and update our internal models^[15]^. In any kind of closed-loop control environment, sensory feedback delays have to be minimized to improve control stability^[16]^: if the feedback arrives too late, it might prove irrelevant to the momentary movement and the ability to update our internal models will be lost due to the inability to establish an accurate causal relationship between performed action and perceived feedback.

The speed of feedback is particularly important when we consider the mechanisms of the decision making process inside the brain^[17,18]^. When a decision is required, the brain gathers so-called evidence in the form of incoming sensory information^[19]^. In the case of movement, we mainly rely on the modalities of vision, touch and proprioception to provide us with information regarding internal and external states to ensure fluid and accurate movement^[12]^. These sources are integrated by weighting them based on the certainty of the information; enough evidence has been gathered when a certain threshold is passed, resulting in a decision^[20]^. The speed and accuracy of our decision depends on the level of this decision-threshold: if the threshold is low, less evidence is needed to make a decision, which speeds up the process while sacrificing accuracy. On the other hand, a higher threshold requires more evidence, which raises the likelihood of a correct answer, but makes the process slower. Due to the decision making process needing time to accumulate evidence necessary to make a judgment, the sensorimotor system is faced with a dilemma: it can make a fast decision which compromises the response accuracy or it can wait for more evidence, compromising the reaction time. In order to minimize the effect of this tradeoff on real-time control, it is not enough to provide sensory feedback through a fast modality, but the feedback also needs to include all the necessary information within a short timescale.

The speed of processing and the impact of the incoming information are different across modalities. These differences are reflected in reaction times, and are dependent on many factors, such as attention, conduction time and complexity of stimuli^[21]^. However, generally, reactions to tactile stimuli are faster than to visual stimuli^[22–24]^. When integrating information from different modalities, how much information is taken from a specific modality is related to the weighting given during sensory integration. The weighting depends on the certainty of the signal, where a modality with high certainty will be weighted higher than those with low certainty^[25]^. As a result, adding feedback is not always going to improve performance: if the supplementary feedback carries a high amount of uncertainty, sensory integration will be inefficient and the impact in control will be negligible. Thus, to be effective, supplementary feedback has to be fast and distinguishable in order to be assigned higher weighting.

In the case of individuals with limb-difference, the tactile and proprioceptive modality are almost completely lost, resulting in difficulties for people to localize their limbs in space, as well as losing the ability to accurately sense their prosthetic limbs interacting with the environment. To compensate, people rely on vision and, in some cases, incidental feedback such as auditory cues^[26]^. As visual stimuli are generally processed more slowly than tactile ones^[22]^, this potentially introduces a significant control delay to the system. This delay is further increased in the case of myoelectric devices due to how EMG signals are usually smoothed via windowing methods^[27]^, which are dependent on collecting data over a certain amount of time before an output can be generated. Furthermore, the overreliance on vision imposes a heavy cognitive load during prosthetic control, which hinders the use of the device^[28]^. In order to alleviate the load on the visual system and improve feedback control, supplementary feedback through other modalities should be provided.

If we want feedback that integrates well into the sensorimotor loop, we need to provide it in a way that is fast, easily recognizable and distinguishable. In this study, we investigated how transcutaneous electrotactile stimulation can fulfill these requirements. Electrotactile stimulation is an attractive choice due its non-invasive and energy efficient nature^[29]^. Furthermore, it has been used before to close the loop during prosthetic control, either by conveying information regarding sense of touch useful to determine grasping force^[30,31]^ or proprioception which helps keep track of joint angles^[32]^. Because of the importance of speed in sensorimotor control, we first aimed to confirm that electrotactile stimulation registers more quickly than visual stimulation. In a second experiment, we investigated how different stimulation- and environment-related parameters can influence the speed-accuracy trade-off (SAT), with the goal of identifying parameters that lead to both fast and accurate decision-making.

## Methods

Participants provided written informed consent before the start of the experiments, agreeing to take part in the study and their data, once anonymized, to be used and shared. The study was approved by the UCD Human Research Ethics Committee – Sciences (LS-22-46-Dupan).

### Experiment 1: Reaction time

#### Experimental protocol and design

Twenty participants (9 male, 11 female, with mean age ± SD = 29 ± 5) without limb difference took part in the experiment. Participants sat in front of a monitor and provided their responses using a game controller (CH Products Flight Stick Pro 200-503). A pair of electrodes (Axelgaard PALS 5×5 flex electrodes, USA) were positioned on the anterior side of the non-dominant forearm of the participants, slightly above the wrist and along the flexor carpi radialis. Electrotactile stimulation was provided using an analog stimulus isolator (A-M Systems Model 2200, USA) which was controlled by an Arduino Uno microcontroller connected to the experimental PC running a Python script. Stimulation was provided as monophasic square waves, with frequency and amplitude fixed at 50Hz and 3mA respectively. These values were chosen due to physiological factors, as well as adherence to existing literature^[30,31,33]^. The perceived stimulus intensity was modulated by increasing and decreasing the pulse width of stimulation trains.

The experiment consisted of 2 parts, which are represented in Fig. 1a. First, the detection threshold (DT) was determined by increasing the stimulation pulse width from 50μs in 50μs increments, with the stimulus trains lasting 5 seconds. Participants were asked to press a button on the controller when they felt the stimulation, which resulted in the pulse width dropping by 100μs. The DT was defined as the intensity at which the participant indicated a perception event 3 consecutive times.

**Figure 1.**
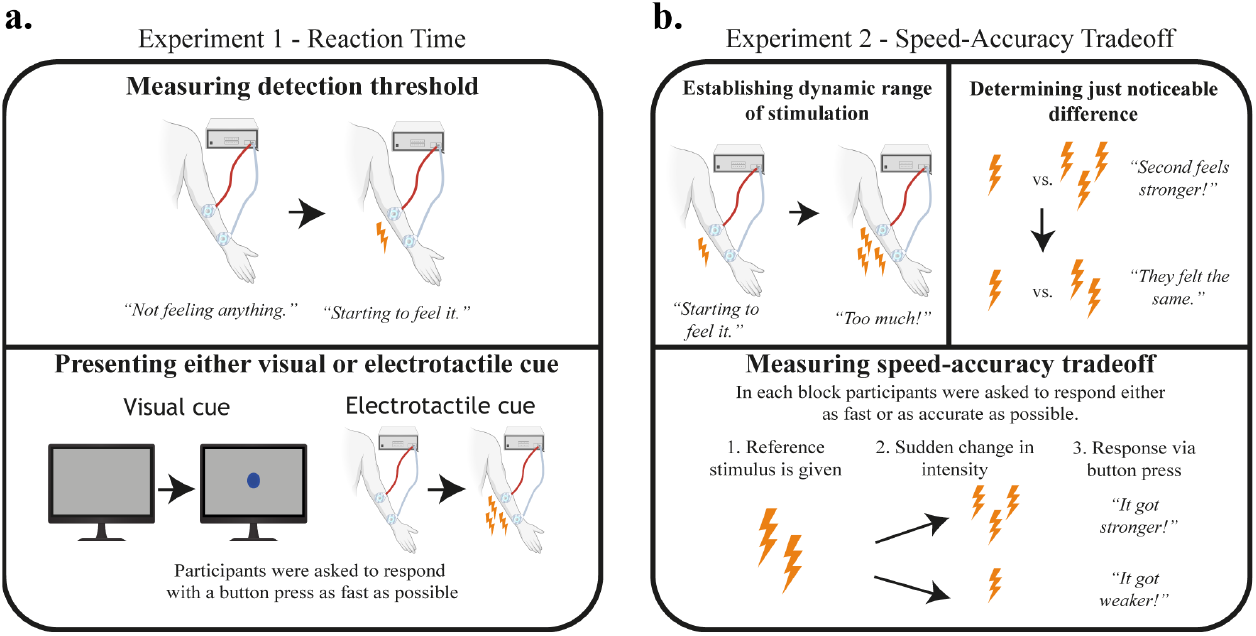
Experimental methods. a: Reaction time experiment methodology. Subject is randomly presented with either a visual or an electrotactile stimulus. They have to respond with a button press as fast as possible. No discrimination between types of stimuli have taken place. b: Speed-Accuracy Tradeoff experiment methodology. At the start, the dynamic range (DR) is established via measuring the detection threshold (DT) and pain threshold (PT). Then participants are presented with two consecutive stimuli with a small delay inbetween to establish the just noticeable difference (JND). Finally, participants indicated if the first or second stimulus felt stronger after an instantaneous fall or rise in intensity.

During the reaction time task participants were asked to press the button on the game controller as fast as possible after perceiving a stimulus. Stimuli were either a visual stimulus (a circular dot appearing on the monitor) or an electrotactile stimulus. The intensity of the electrotactile stimulus was set to twice the value of the DT. The experiment consisted of 2 blocks with 40 trials each. The order of the visual and electrotactile stimuli were randomized on the level of individual trials. The start of each trial was indicated by a cross-shaped visual stimulus appearing on the screen, while the end of trial was denoted by the cross surrounded by a color-coded circle (blue for visual and orange for electrotactile trials), which indicated a received response. The delay between the start of the trial and stimulus onset were randomized in an interval of 2-4 seconds in order to avoid the participants predicting the time of stimulus onset. The stimuli in both conditions were deemed to be easily noticeable in pilot trials, thus a timeout condition was not established.

#### Data analysis

The main outcome variable used to determine the difference in processing speed between visual and electrotactile stimulation was reaction time. For each feedback modality, average and standard deviation of reaction times were determined for each participant. These values were used for subsequent analysis. Normality was tested using the Shapiro-Wilk method which confirmed normal distribution of the averages, but not for the standard deviations. Therefore, we used the Wilcoxon signed rank test for all comparisons. Furthermore, we fit an exponentially modified gaussian distribution to the pooled reaction times from all participants. This was done in order to confirm that both sets of reaction times conform to established distribution patterns^[34,35]^. For the fitting of the exponentially modified gaussian, we used the *Exponentially modified Gaussian (ex-Gaussian) distributions toolbox* available through MATLAB File Exchange^[36]^.

### Experiment 2: Speed-Accuracy Tradeoff (SAT)

#### Experimental protocol and design

Twenty participants (11 male, 9 female with mean age ± SD = 28 ± 5) without limb difference took part in the experiment. Participants sat in front of a monitor and provided their responses using a numeric keypad. Similarly to the first experiment, the electrotactile stimulation was provided using an analog stimulus isolator (A-M Systems Model 2200). The stimulus isolator was controlled using an NI board (NI USB-6211) connected to the experimental PC running a Python script. Stimulator electrodes were placed above the flexor carpi radialis of the non-dominant forearm of the participants. Stimulation was provided as biphasic square waves, with frequency and amplitude fixed at 50Hz and 5mA respectively. Due to the higher amplitude of stimulation enabled by the NI board, the stimulation electrodes were placed along the upper part of the flexor carpi radialis; this elicited a much more defined sensation compared to the 3mA stimulation provided by the Arduino board. Perceived stimulus intensity was modulated by increasing and decreasing the pulse width.

Fig. 1b shows the 3 parts of this experiment. First, we measured the dynamic range (DR) of stimulation via establishing the detection threshold (DT) and pain threshold (PT). The DT was determined the same way as in the reaction time experiment. Following that, the pulse width was increased to a point where the participant reported a feeling of pain or discomfort. For the comfort of the participants, this level was only recorded once, and was noted as the pain threshold (PT). The DR was determined as the pulse width range between the DT to the PT.

In the second part, the just noticeable difference (JND) was measured with respect to a stimulus set to 25% of the DR using a staircase procedure. The procedure began with the participant receiving two consecutive stimuli with a 0.5s pause in between each; one stimuli was set to 25% of the DR (reference stimulus) and the other set to 50% of the DR (compared stimulus). In each of the trials, the order of the reference and compared stimuli were randomized. The participant was instructed to determine the stronger of two consecutive stimuli after receiving both. Every correct response resulted in the pulse width of the compared stimulus dropping by 10μs and every incorrect response in the pulse width rising by 20μs. An incorrect response followed by a correct one, or a correct response followed by an incorrect one were defined as reversals. JND was determined as the difference between the reference intensity and the average of the ten intensities where a reversal has taken place.

During the main SAT task, participants were asked to respond to an instantaneous shift in stimulus intensity via a button press, indicating whether the reference or shifted stimulus felt stronger. The instantaneous shift aimed to replicate the provision of electrotactile feedback during continuous prosthetic control. The reference intensity was set at 25% DR and the magnitude of change in intensity corresponded to 1-4 times JND. The direction of intensity shift (rising/falling) was randomized over trials. The first stimulus lasted between 2 and 4 seconds, while the second stimulus continued until the participants responded with the button press or timed out after 3 seconds. At the start of each block, participants were instructed to prioritize speed or accuracy in their response. In half of the trials, cognitive load was imposed during the task. This was done in order to investigate how the increased cognitive load present during prosthesis use^[28]^ affects the speed and accuracy of recognizing changes in electrotactile stimulation.

To impose cognitive load, participants were asked to simultaneously listen to paragraphs from an audiobook (Dr. Ox’s Experiment by Jules Verne) and answer multiple choice questions (MCQ) afterwards in half of the blocks^[37–39]^. Speed/Accuracy conditions were changed every block, while cognitive load conditions were changed every two blocks. The outcome variables of the task were response speed and response accuracy.

#### Data analysis

The main outcome variables used to determine the impact of stimulation parameters in instantaneously shifting electrotactile stimuli were reaction time and response accuracy. The independent variables we considered were: (1) stimulus shift magnitude, (2) stimulus shift direction, (3) presence of added cognitive load and (4) explicit instruction regarding focus on speed or accuracy of response.

An ANOVA test was performed to determine which stimulation parameters had an effect and whether interactions were present between them for both reaction times and response accuracies. Only the per-condition average reaction times and response accuracies were considered for each participant. Normality was tested using the Shapiro-Wilk method for all subsets of the data which confirmed normality in 13 of 20 subsets. Even though an ANOVA was used to determine interactions and individual effect, due to the high amount of non-normal subsets, we used the Wilcoxon signed rank test to compare conditions. In the case of shift magnitude comparisons, Bonferroni correction was applied to the p-values of Wilcoxon signed rank tests.

## Results

### Experiment 1: Reaction time

We first tested whether there is a significant difference in reaction times between visual and electrotactile stimulation. Based on existing literature stating that tactile stimulation registers faster than visual stimulation in general^[22–24]^, our tests aimed to confirm whether this is true for electrotactile stimulation as well. A Wilcoxon signed rank test revealed that average reaction times are significantly faster for electrotactile stimulation than visual stimulation (p=0.008). Comparing the standard deviations of the RTs for both modalities revealed no significant difference (p=0.067), while taking the averages of the 10th, 50th and 90th percentiles of each participant’s reaction times showed significantly lower reaction times in all three percentiles (Fig. 2a). This suggests that electrotactile stimulation doesn’t show significantly increased variability compared to visual stimulation while allowing faster reactions on average. Fig. 2b shows the exponentially modified gaussian which was fitted to the pooled reaction times of both stimulus modalities^[34,35]^. The fitted distributions confirm that there is a 56ms difference between the means of the distributions, with the electrotactile modality being faster (μ_VIS_=299.6ms and μ_ELE_=243.2ms)

**Figure 2.**
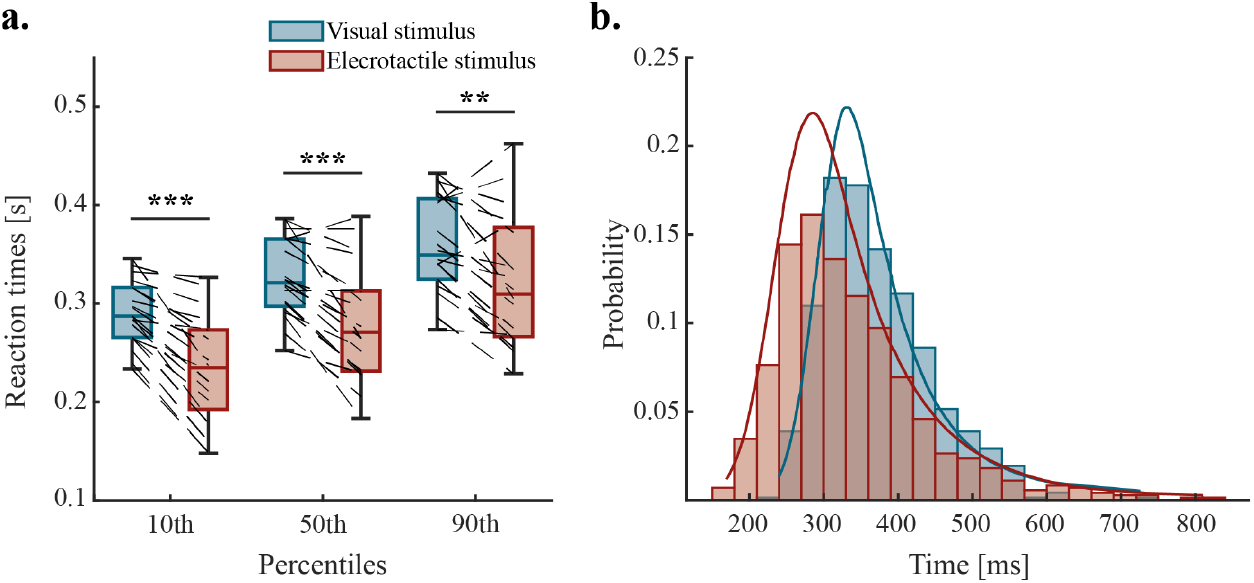
Results of the reaction time experiment. a: Average reaction times are significantly lower for electrotactile stimulation in the 10th, 50th and 90th percentiles (p<0.001, p<0.001 and p=0.0032 respectively). Connected data points signify the percentile averages for individual participants; b: Distribution of pooled reaction times. Reaction times to both stimuli follows an exponentially modified gaussian distribution.

### Experiment 2: SAT

The second experiment investigated whether the SAT can be influenced by modulating stimulus-related (stimulus shift direction and magnitude) and environmental variables (explicit instructions and presence of cognitive load) during shifting electrotactile stimuli. Identifying the parameters that ensure fast and accurate perception of electrotactile stimuli is a prerequisite for implementation of electrotactile feedback in prosthetic control. The experiment was aimed to measure reaction times and correct responses to instantaneous shifts in electrotactile stimulation under four different sub-conditions: stimulus shift direction (rising and falling intensity), stimulus shift magnitude (1-4xJND value), explicit instruction (focus on speed or accuracy of response) and concurrent cognitive load (audio on and off).

Four-way ANOVA was performed on both reaction time and response accuracy data in order to determine which variables had the most effect on the outcomes, as well as to find interaction effects between them. The ANOVA revealed that stimulus shift magnitude had the strongest effect on both response accuracies and reaction times (p<0.001 in both cases). Significant differences can be seen for all intensities in response accuracies, except for 3x and 4xJND values (p<0.001 for all 1xJND comparisons, p = 0.0032 for comparison between 2x and 3xJND). Pairwise comparisons show significant differences across all intensities for response times (p<0.001 in all cases).

Most response inaccuracies were observed at the lowest shift magnitude. At 1xJND 43.38% of all answers were incorrect answers; this includes missed responses which constituted 35.96% of all answers at the lowest shift magnitude (Fig 3a.). The number of missed and inaccurate responses decreased significantly as shift magnitude was increased (Fig. 3a-b). In terms of accuracy, median response accuracy at 1xJND was 61.67%, which significantly increased to 91.67% at 2xJND level. Response accuracies further increased at the 3xJND and 4xJND levels to 96.25% and 96.67% respectively. A similar effect was observed in the case of reaction times for correct responses, which decreased with increasing shifts (Fig. 3c): median correct response times at 1xJND level were 1.439 seconds. Each decrease in the shift reduced median response times, with 4xJND being the lowest at 0.691 seconds, showing an almost 750 millisecond drop in response time from 1x to 4xJND. These results imply that providing larger shifts in electrotactile stimulation improves discriminability and results in faster and more accurate perception.

**Figure 3.**
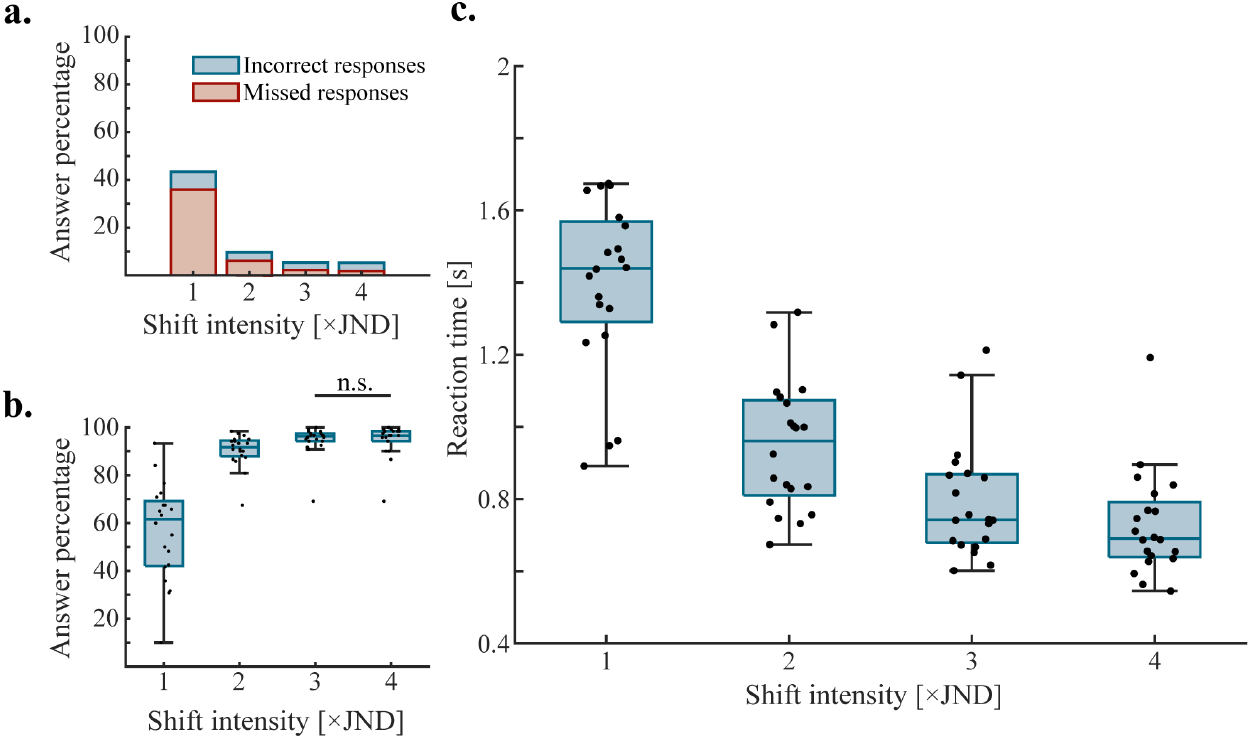
Response accuracies and reaction times as a function of shift intensity; a: The number of incorrect and missed responses decrease as shift intensity gets larger. b: Average response accuracy per participant increases with shift intensity. Black dots denote averages of individual participants. Only non-significant differences are shown, all other comparisons are significantly different. c:Average reaction times per participant decrease with shift intensity; only correct responses were considered in this case. Black dots denote averages of individual participants. All comparisons were significantly different.

Performing the task with simultaneous cognitive load led to significantly higher reaction times (p<0.001, Fig. 4a), with a median difference of 175ms. While the average number of accurate responses were not affected (p=0.420), there was a significantly higher number of inaccurate responses in the cognitive load condition, not counting missed responses (p = 0.012, Fig. 4b). After blocks with the cognitive load condition, participants were asked to answer questions regarding the audiobook in the form of a MCQ test. Participants on average scored 79% ± 20.5% on the MCQ test, indicating that participants performed well on the secondary task despite their divided focus. In the case of shift direction (rising/falling intensity), the ANOVA revealed an interaction effect with shift magnitude regarding response accuracies (p<0.001). Further analysis using a Wilcoxon signed rank test revealed that a significant difference was only present for the lowest shift magnitude (p=0.0051, Fig. 4c), making rising shifts harder to determine correctly. Explicit instructions regarding focus on response speed or response accuracy did not have a significant effect on either reaction times or response accuracies (p=0.1086 and p=0.2476, respectively). Further Bayes factor analysis confirmed these results, revealing BF_10_ = 0.2285 and BF_10_ = 0.2331 for reaction times and response accuracies.

**Figure 4.**
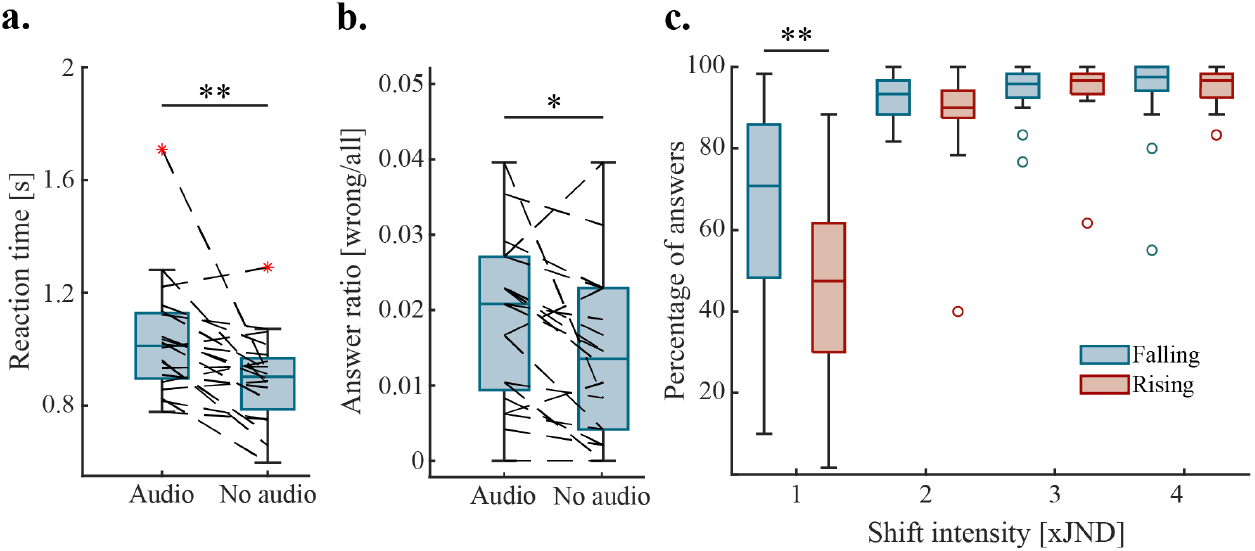
Effects of cognitive load and shift direction on reaction times and response accuracy; a: Added cognitive load results in increased average reaction times; response accuracies do not change significantly. b: Added cognitive load results in more incorrect responses; missed responses are not included and are unaffected (one outlier not pictured) c: There are less correct responses on average for rising shift than falling shift for the smallest shift intensity; there is no significant difference at higher shift intensities.

## Discussion

Using a prosthetic device effectively requires its incorporation into the sensorimotor system. If supplementary feedback is added to the prosthetic device, its parameters need to be designed to provide fast and distinguishable information which the sensorimotor system can exploit. In this study, we showed that (1) bigger steps during shifting electrotactile stimulation lead to faster and more accurate responses and (2) the presence of cognitive load only increases response times to shifting electrotactile stimulation, but does not affect the number of correct responses. Furthermore, we have confirmed previous findings which demonstrate that reaction times to electrotactile stimuli are faster than those to visual stimuli. These results reinforce the notion that providing bigger steps during electrotactile stimulation lessen the effect of the SAT and make the perception of shifting stimuli easier. Therefore, providing electrotactile stimulation in bigger steps could improve closed-loop control of prosthetics by making stimuli faster and less cognitively demanding to respond to.

Current research efforts focusing on providing supplementary feedback for upper-limb prosthetics aim to improve control via encoding tactile or proprioceptive events^[30–32]^. In order to improve control, we have to ensure the feedback is properly integrated into the sensorimotor control loop^[12]^. Proper integration can be facilitated by providing feedback through a fast modality. While the advantage in reaction times for the tactile modality is promising, it is important to remember that sensory stimuli carry varied information and the process of distinguishing them takes time as well^[17,18]^. In order to avoid delays and inaccuracies, the changes in sensory feedback need not only to be swiftly noticeable, but easily discernible as well. Our results show that introducing bigger steps in electrotactile stimulation intensity leads to positive effects for both response times and response accuracy. The effect of step size on the response times is interesting in the context of previous research, which reported that modulation of perceived intensity in small increments (1xJND) during a closed-loop tracking task was characterized by a noticeable control delay^[40,41]^. Our results suggest that while small increments enable a large number of levels of stimulation within the DR, they also increase the time it takes to accurately process the sensation of shifted stimulation intensity. Thus, we argue that providing electrotactile stimulation using bigger steps in perceived intensity could result in faster, and therefore more reliable closed-loop control for prosthetics.

Prosthesis users experience heavy cognitive load arising from several sources, such as overreliance on the visual system^[42]^ and the undergoing reorganization of the sensorimotor system^[43]^. Additionally, everyday situations require users to manipulate their device while performing a simultaneous task (eg. holding a conversation while using the prosthesis) which further increases the amount of cognitive load. To simulate these circumstances, we introduced a cognitive load condition into the experiment. While our results imply that presence of cognitive load has no effect on the number of accurate responses, it is important to note that it results in a significant increase in response times and incorrect answers. The increase in response times may lead to control delays in a closed-loop control task, which is in line with existing research which has shown that presence of cognitive load slowed down movement during a closed-loop dual task for both invasive and non-invasive feedback^[46]^. However, a different study found no impairment in control performance in the presence of cognitive load^[44]^. This discrepancy could be the result of a difference in task nature, as the ‘box and blocks’ task requires less rapid response to incoming stimuli compared to our task or the walking task.

During movement, choosing the appropriate motor command is dependent on incoming stimuli, which are subject to continuous change; this often results in us having to alter or adapt our motor commands as well^[18]^. When presented with a stimulus which requires a change in our motor command, we have to make a decision on how to alter our movement. This decision, ideally, is both fast and accurate in assessment of the change necessary; in reality, the speed-accuracy tradeoff in most situations prevents us from fulfilling both conditions at the same time. One of the most common ways to manipulate the SAT is through verbal instructions, as in explicitly instructing the participant whether to focus on speed or accuracy in their response^[45–47]^. While we implemented verbal instructions in our second experiment, we have failed to see significant differences in this condition regarding both response times and response accuracies, which could be attributed to the qualitative nature of verbal instructions in SAT tasks^[20]^. Another possible explanation for the similar performance in the ‘speed’ and ‘accuracy’ conditions is related to task difficulty: harder tasks tend to have higher response times and reduced accuracy, while in easier tasks response times decrease and response accuracies increase^[48]^. Although our results show low response accuracies and high response times in the 1xJND condition, both of these metrics improve significantly for the 2xJND condition. While we see further improvements regarding response times in the 3x and 4xJND conditions, average response accuracies already reach near 100% average accuracy for the 3xJND condition with a non-significant difference in the 4xJND conditions. This behavior is reminiscent of the tail ends of the psychometric and chronometric curves, with 1xJND representing the beginning and 2xJND already showing signs of the opposite tail end. This implies that the discrimination threshold - where we would see a difference in the verbal instruction condition - could be between these two conditions^[49,50]^.

During use of a prosthetic device, the user heavily relies on their vision in order to gain feedback from the device. While vision provides reliable feedback, it lacks the advantage in speed the tactile modality possesses^[21,22,24]^. Together with the increase in cognitive load, this insufficiency in the speed of the visual modality introduces further delay into the sensorimotor control loop, further destabilizing the system. For this reason, supplementary feedback should be provided through a modality that is ideally faster than the visual modality. Although it has been shown before that tactile stimuli are generally registered faster than visual stimuli^[51]^, this has not yet been confirmed for electrotactile stimulation. In order to test this, we performed a reaction time test where we compared average reaction times for both visual and electrotactile stimulation. Our results showed that average response times to electrotactile stimulation were ~50ms faster than visual stimuli, which confirmed our hypothesis. Moreover, this difference was already observed in the averages of the fastest 10% of participants’ reaction times between visual and electrotactile stimulation. The difference between reaction times already appearing in the leading edge of the reaction time distribution suggests that the advantage of electrotactile stimulation is due to the faster encoding of evidence rather than necessarily a change in the evidence accumulation process itself. Shifts in the leading edge of RT distributions in perceptual decisions are best explained by a difference in the non-decision time parameter of an evidence accumulation model, which encompasses both sensory encoding and motor processes^[52]^. The difference in sensory encoding time is thought to arise from differences in the mechanisms of the peripheral nervous system across sensory modalities, leading to variations in their associated reaction times^[53]^.

The main limitation of our study was that testing was performed on able-bodied participants. However, the ability to distinguish between different intensities of electrical stimulation during a sensorimotor task were shown to be similar between able-bodied people and people living with limb-difference^[54]^. It is important to note that tasks included in this research were feed-forward in nature. While it is true that closed-loop tasks more closely reflect the control of a prosthetic device, these studies were necessary to determine the delays associated with different feedback modalities and their stimulation parameters. This study allowed us to identify important stimulation parameters for future testing in closed-loop control. Lastly, we used an ANOVA to determine possible interactions and the influence of parameters at group level. However, not all subsets of the data were normally distributed. This decision was made due to the lack of suitable alternatives to ANOVA for non-normally distributed datasets. All subsequent comparisons between subsets were performed via methods suited for non-normally distributed data.

## Conclusions

In this study, we explored how to optimize electrotactile stimuli for prosthetic feedback to be integrated into the sensorimotor system. We have shown that bigger steps during shifting electrotactile stimulation lead to faster and more accurate responses. Our results also show that presence of cognitive load only increases response times to shifting electrotactile stimulation, but does not worsen response accuracy. Therefore, providing electrotactile stimulation in bigger steps could improve closed-loop control of prosthetics by making electrotactile stimuli easier and less cognitively demanding to respond to.

## Declarations

### Ethics approval and consent to participate

Participants provided written informed consent before the start of the experiment, agreeing to take part in the study and their data, once anonymized, to be used and shared. The study was approved by the UCD Human Research Ethics Committee – Sciences (LS-22-46-Dupan).

### Consent for publication

Not applicable.

### Availability of data and materials

The datasets used and analyzed during the current study are available from the corresponding author on reasonable request.

### Authors’ contributions

FJ designed and carried out the experiments, analyzed the data and drafted the manuscript. EC was involved in the design of the SAT experiment and analysis, interpretation of the data, and revision of the manuscript. SD was involved in conception and design of the study, carried out interpretation of the data and revision of the manuscript. All authors have read and approved the manuscript.

## Acknowledgements

The authors would like to acknowledge the help of Prof. Simon Kelly for his insightful comments on experimental design and statistical analysis.

